# Overcoming resistance to KRAS inhibition in KRAS/PIK3CA co-mutated colorectal cancer

**DOI:** 10.64898/2026.11.10.765758

**Authors:** L. Ferret, J.L. Montaño, K.B. Nayak, R.E. Turnham, C.G. Rivera, R. Xu, P. Kongpracha, D.L. Swaney, N.J. Krogan, J.D. Gordan

## Abstract

Oncogenic alleles of *KRAS* drive colorectal cancers (CRC) growth and are associated with worse disease outcomes. Despite potent allele-specific and pan-RAS inhibitors targeting now in clinical test, CRC patients are not experiencing the same therapeutic benefit as patients with other cancers. PI3-Kinase (PI3K) is a direct RAS effector and is co-mutated in about 45% of CRC cases, raising the possibility that frequent oncogenic *PI3K* mutations contribute to the limited benefit of KRAS inhibition in CRC. Using isogenic colorectal cancer cells engineered to harbor either of the two most common activating *PIK3CA* mutations, we show that oncogenic PI3K promotes resistance to the pan-RAS inhibitor daraxonrasib (RMC-6236). Global phosphoproteomic profiling revealed that *PIK3CA*-mutant cells undergo a distinct adaptive response to RAS inhibition, characterized by sustained PI3K-AKT signaling and extensive remodeling of pathways controlling cell-cell adhesion and junction signaling, alongside suppression of apoptotic signaling. Combined RAS and PI3Kα inhibition overcame this resistance, suppressed signaling through both pathways and induced apoptosis in *PIK3CA*-mutant cells. Together, these findings identify oncogenic *PIK3CA* as a determinant of resistance to RAS inhibition in *KRAS*-mutant colorectal cancer and provide a mechanistic rationale for combined KRAS and PI3Kα inhibition in *KRAS*/*PIK3CA* co-mutant tumors.

## INTRODUCTION

RAS proteins are among the most frequently altered oncogenic drivers in human cancer. KRAS accounts for more than 80% of mutated *RAS* in solid tumors, and is a major driver of pancreatic, lung, and colorectal cancers (CRC)^1, 2^. In CRC, *KRAS* mutations are found in approximately 40% of tumors, predominantly affecting codon 12, with G12D and G12V being the most prevalent variants^3^.

KRAS is a membrane-associated small GTPase that cycles between an active, GTP-bound (“ON”) state and an inactive, GDP-bound (“OFF”) state. Oncogenic *KRAS* mutations impair this regulatory cycle and favor the accumulation of the active, GTP-bound protein, resulting in sustained activation of downstream signaling pathways, including the RAF-MEK-ERK and PI3K-AKT axes^4^. Daraxonrasib (RMC-6236; DRX) is a pan-RAS(ON) inhibitor that targets the GTP-bound state of multiple RAS variants. Daraxonrasib has demonstrated clinical activity across RAS-mutant tumors and was recently FDA approved for metastatic pancreatic adenocarcinoma^5^. However, its activity and the molecular determinants of response in *KRAS*-mutant CRC remain poorly understood.

This question may be particularly relevant in CRC as the consequences of KRAS inhibition are shaped not only by adaptive upstream signaling, exemplified by the addition of EGFR antibodies (cetuximab, panitumumab) to KRAS G12C inhibitors to counteract RTK-driven pathway reactivation ^6, 7^ , but also by parallel downstream survival pathways that can maintain tumor growth despite the suppression of RAS-MAPK signaling. Among these, the PI3K-AKT pathway represents a major candidate determinant of response and resistance^8^.

PI3K signaling constitutes a major downstream effector arm of oncogenic RAS. Despite the typical mutual exclusivity of *KRAS* mutations with those of its downstream effectors (e.g. *BRAF*), activating mutations in *PIK3CA*, encoding the p110α catalytic subunit of PI3K, co-occur with approximately 40% of KRAS mutations in CRC,^9^ with H1047R and E545K representing the predominant hotspot mutations in the kinase and helical domains, respectively. Importantly, extensive crosstalk and feedback exist between these pathways as inhibition of one signaling pathway can promote the compensatory activation of the other, providing a mechanism for treatment resistance and limiting the efficacy of single-pathway inhibition^8^. These observations raise the possibility that PIK3CA co-mutation and persistent PI3K–AKT signaling may represent important determinants of response to RAS(ON) multi-selective inhibition in KRAS-mutant CRC and provide a rationale for therapeutic strategies that simultaneously suppress both signaling axes.

Here, we show that oncogenic *PIK3CA* mutations (H1047R and E545K) promote resistance to pan-RAS inhibition in *KRAS*-mutant CRC. Additionally, in global phosphoproteomics data we find that the mechanistic basis for the observed resistance to KRAS inhibition is mediated by sustained PI3K-AKT signaling, apoptosis inhibition and remodeling of cell-cell adhesion and junction signaling in KRAS/PIK3CA co-mutated CRC models. Most importantly, concomitant KRAS and PI3Kα inhibition effectively overcomes resistance to KRAS inhibition by driving co-mutated CRC cells towards apoptosis. Co-targeting of PI3K and MAPK pathways had previously been precluded by the toxicity of compounds targeting mutant and WT variants of the same protein. However, with the emergence of allele-specific KRAS and PIK3CA targeting agents, selectively targeting mutants in both pathways is an intriguing path forward in CRC therapy that may overcome some of the barriers of targeting KRAS alone.

## MATERIALS AND METHODS

### ANTIBODIES AND REAGENTS

#### Reagents

- Daraxonrasib (RMC-6236) (MedChemExpress, HY-148439)
- Inavolisib (GDC-0077) (MedChemExpress, HY-101562)
- Alpelisib (MedChemExpress, HY-15244)
- SYTOX-green (ThermoFisher Scientific, S7020)
- DMSO (Sigma-Aldrich, D2650)

#### Antibodies

- PIK3CA rabbit (Cell Signaling Technology, 4249)
- Flag mouse (Sigma-Aldrich, A8592)
- β-Actin mouse (Cell Signaling Technology, 3700)
- Phospho-AKT S473 XP rabbit (Cell Signaling Technology, 4060)
- AKT mouse (Cell Signaling Technology, 2966)
- Phospho-MEK1/2 S2217/221 rabbit (Cell Signaling Technology, 9154)
- MEK1/2 rabbit (Cell Signaling Technology, 9122)
- Phospho-S6 Ser235/236 rabbit (Cell Signaling Technology, 4857)
- S6 rabbit (Cell Signaling Technology, 2217)
- Phospho-ERK1/2 Thr202/Tyr204 rabbit (Cell Signaling Technology, 4370)
- ERK1/2 rabbit (Cell Signaling Technology, 4695)
- Vinculin XP rabbit (Cell Signaling Technology, 13901)
- Cleaved PARP rabbit (Cell Signaling Technology, 5625)
- Cleaved Caspase-3 rabbit (Cell Signaling Technology, 9664)

### CELL CULTURE

The SW837 (human colorectal adenocarcinoma) and SW620 (human colorectal adenocarcinoma) cell lines were purchased from the American Type Culture Collection (respectively, CCL-235™, CCL-227 ™). GP2d (human colorectal adenocarcinoma) cell line was obtained from Merck (Sigma-Aldrich, 95090714). T84 (human colorectal carcinoma) cell line was obtained from UCSF biobank.

SW837 and SW620 cell lines were all cultured in Leibovitz’s (L-15; Thermo Fisher Scientific) culture medium supplemented with 10% fetal bovine serum (Life Technologies) and 1% PenStrep (Gibco), at 37C in a humidified incubator without CO2. T84 were grown in DMEM/F-12 (Thermo Fisher Scientific, 11320033), GP2d cells were grown in Dulbecco’s Modified Eagle’s (DMEM, Corning) culture medium supplemented with 10% fetal bovine serum and % PenStrep, cultivated at 37C in a humidified incubator with 5% CO2. All the cell lines were free from myco-plasma contamination (MycoAlert® PLUS Mycoplasma Detection Kit, Lonza LT07-710).

### CRISPR/CAS9-ENGINEERED ISOGENIC *KRAS*/*PIK3CA* CO-MUTATED MODELS

CRISPR–Cas9-mediated homology-directed repair (HDR) was used to generate cells carrying the H1047R/E545K PIK3CA mutation. A synthetic H1047R or E545K-targeting guide RNA (gRNA; Integrated DNA Technologies) was complexed with Cas9 nuclease (IDT) in DPBS (Corning) to form a ribonucleoprotein (RNP) complex containing 4.8 μM gRNA and 4 μM Cas9 final concentrations. RNP complexes were incubated for 20 min at room temperature before nucleofection.

For each reaction, 200,000 cells were collected by centrifugation at 200G for 5 min. Cell pellets were resuspended in 20 μL freshly prepared supplemented P3 nucleofection solution, consisting of 16.4 μL of P3 Primary Cell 4D-Nucleofector X Solution and 3.6 μL Supplement (Lonza, V4XP-3032).

For HDR-mediated introduction of the H1047R or E545K mutation, an HDR donor oligonucleotide (IDT) containing the desired nucleotide substitution was used as the repair template. The nucleofection reaction was assembled in a final volume of 30 μL and contained 200,000 cells in 20 μL supplemented P3 solution, 5 μL of the Cas9–gRNA RNP complex, 4 μM of H1047R HDR donor oligonucleotide (IDT), 1.2 μL Alt-R Cas9 Electroporation Enhancer (IDT), and 2.6 μL DPBS (Corning).

This 30-μL reaction was transferred to a nucleofection strip and electroporated using a 4D-Nucleofector system (Lonza) with the pulse program appropriate for the cells. Immediately after nucleofection, 75 μL of pre-warmed complete growth medium was added to each nucleofection well. Cells were gently resuspended and transferred to a 24-well plate containing 1 mL of complete growth medium supplemented with 1.7 μL Alt-R HDR Enhancer V2 (IDT) per well.

After 24h, the medium was replaced with fresh complete growth medium without HDR Enhancer V2. Following recovery and expansion, cells were divided between two wells of a 6-well plate: one population was maintained for further culture, while the other was used for genomic DNA extraction using GeneJET Genomic DNA Purification Kit (ThermoFisher Scientific, K0722). The targeted genomic region encompassing the H1047R or E545K editing site was amplified by PCR using Q5® Hot Start High-Fidelity DNA Polymerase (NEB, M0493S). PCR products were subsequently analyzed by Sanger sequencing to determine the genotype of individual clones and classify them as unedited, heterozygous, or homozygous for the H1047R or E545K mutation.

### TET-ON INDUCIBLE FLAG-PIK3CA CO-MUTATED MODELS

Doxycycline-inducible plasmids encoding FLAG-tagged PIK3CA WT, H1047R, or E545K were kindly provided by the N. Krogan laboratory. Lentiviral particles were produced in HEK-293T cells by co-transfection of the respective PIK3CA expression plasmid (4 μg) with the GPTR packaging plasmid (1.3 μg) and VSV-G envelope plasmid (0.7 μg). Plasmid DNA was mixed with 14 μL FuGENE® 6 transfection reagent (Promega, E2692) in 100 μL Opti-MEM™ (Thermofisher Scientific, 31985062) and the transfection mixture was added dropwise to HEK-293T cells cultured in 6-cm dishes. Viral supernatants were collected 48–72 h post-transfection, centrifuged at 500 × g to remove cellular debris, and filtered through a 0.45-μm filter.

SW837 and SW620 cells were transduced with 1 mL of viral supernatant per well of a 6-well plate in the presence of 8 μg/mL polybrene (EMD Millipore, TR-1003-G). After 24 h, the viral supernatant was replaced with fresh complete growth medium. Stably transduced cells were selected with puromycin (1 μg/mL) for 3–5 days.

Expression of the FLAG-tagged PIK3CA constructs was induced with doxycycline (1 μg/mL). Successful generation of the inducible cell lines and doxycycline-dependent expression of the constructs were confirmed by Western blot analysis using antibodies against FLAG and PIK3CA, comparing doxycycline-treated and untreated conditions.

### CELL VIABILITY ASSAY

The Cell Titer Glo assay was used to monitor cell viability after drug treatment. Cells were seeded in a white 96-well plate (4,000 cells/well) in triplicates. The following day, cells were treated with the indicated drugs for 5 days. After 5 days of treatment, reconstituted CellTiter-Glo® Reagent (Promega, G7571) was added 1:5 in DPBS (Corning) to each well and incubated for 10 min. The absorbance in each well was measured at 570 nm using an absorbance plate reader (BioTek Synergy 2). The IC50 of the drugs was determined using a non-linear regression curve within GraphPad Prism V10.1.1 software.

### IMMUNOBLOTTING

Adherent cells were rinsed with ice-cold DPBS (Corning), collected by scraping and lysed in RIPA lysis buffer (Pierce) with complete protease inhibitor (Sigma Aldrich) and phosphatase inhibitor (Roche). Samples were incubated on ice for 20 min with intermittent mixing and subsequently centrifuged at 14,000 × g for 10 min at 4°C. Clarified cell lysates were quantified through BCA assay according to the manufacturer’s instructions. Absorbance was measured at 562 nm using a microplate reader (BioTek Synergy 2). Equal amounts of lysate extracts were boiled in 5x loading dye and loaded onto a 4–12% Bis-Tris gel (Invitrogen) and transferred to nitrocellulose membranes (using a standard wet electroblotting system (Bio-Rad)). Membranes were blocked for 1 hour in 5% Albumin bovine fraction V (BSA; Research Products International, A30075) +0.02% sodium azide (Sigma Aldrich, S2002) in TBS-T (Tween20; Sigma-Aldrich, P1379), and proteins of interest were analyzed using primary antibodies (1:1000 in 5% BSA in TBS-T) and corresponding secondary antibodies (Mouse/Rabbit 1:10,000 in 5% BSA in TBS-T).

Immunoreactive proteins were detected using an enhanced chemiluminescence (ECL) substrate (Pierce) and chemiluminescent signals were detected by exposure to X-ray films. Especially for phosphor/total proteins, Restore Plus Stripping Buffer (ThermoFisher Scientific) was used.

### SYTOX-CELL DEATH DETECTION USING LIVE-CELL MICROSCOPY

The quantification of SYTOX-positive cells, indicative of cell death following treatment, was performed using live-cell microscopy. Cells were seeded in black, clear-bottom 96-well plates at a density of 5,000 cells per well. The following day, cells were treated with the indicated drugs in complete medium supplemented with 30 nM SYTOX™ Green reagent (ThermoFisher Scientific, S7020). Immediately after treatment, plates were transferred to an Incucyte® live-cell imaging system (Sartorius) under appropriate culture conditions, with or without CO₂ depending on the cell type. Images were acquired every 4 or 6 h for 5 days, with four images captured per well at each time point.

Brightfield images were acquired using transmitted-light illumination and used to generate cell masks by semantic segmentation using the image analysis software integrated into the Incucyte® system. SYTOX Green fluorescence was detected using the green channel.

For image analysis, SYTOX-positive cells were identified using the image analysis software integrated into the Incucyte® system. Cell objects were segmented from the background based on morphological parameters, including object size. SYTOX-positive cells were defined as detected cell objects exhibiting green fluorescence above a predefined intensity threshold. Cell death was quantified based on the number of SYTOX-positive cells.

### GLOBAL PHOSPHOPROTEOMICS

#### Cell treatment and phosphopeptides enrichment

SW837 CRISPR-Cas9 edited clones unedited (UE2), PIK3CA H1047R (clone 10) or E545K (clone 17) were grown in 15 cm dishes and treated with 100 nM RMC-6236 for 4 h or 24 h, or with an equivalent volume of DMSO. Nine experimental conditions in triplicates resulted in 27 samples processed and acquired independently.

Cells were scraped directly into room-temperature urea lysis buffer (20 mM HEPES pH 8.0, 9 M urea, 1 mM sodium orthovanadate, 2.5 mM sodium pyrophosphate, 1 mM β-glycerophosphate), sonicated and cleared at 20,000 × g for 15 min. Protein input was quantified through a BCA-assay and normalized to 1 mg per sample. Samples were then reduced with dithiothreitol for 30 min at 55C and then alkylated with iodoacetamide for 15 min at room temperature in the dark. Samples were then diluted to a final concentration of 2 M urea using 20 mM HEPES pH 8.0 and then digested overnight with mixing at room temperature with 10 ug of trypsin (Promega, V5280). Digests were acidified to 1% trifluoroacetic acid (TFA), desalted using Sep-Pak C18 columns (Waters, WAT023590) and and dried down in a speedvac vacuum concentrator.

re-suspended in IMAC Fe-NTA loading buffer (0,1% TFA, 85% acetonitrile) and phosphopeptides were enriched using PureCube Fe-NTA magnetic beads (Cube Biotech, 31501-Fe) following the PTMScan Phospho-Enrichment IMAC Fe-NTA Magnetic Beads protocol (CST, 20432). PureCube Fe-NTA magnetic beads were washed (0,1% TFA, 80% acetonitrile) prior to phosphopeptide enrichment. Re-suspended peptides were then added to prepared Fe-NTA beads and binding was performed in 2 ml protein low-binding tubes with rotation for 2h at 4C. Fe-NTA beads were washed with wash buffer and eluted twice using elution buffer (50% acetonitrile, 2.5% ammonia). The eluates were acidified with TFA and dried using a speedvac. Finally, phosphopeptides were re-suspended in a solvent system consisting of mobile phase A (0.1% formic acid in water) and mobile phase B (80% acetonitrile, 0.1% formic acid) and filtered (Millipore, MAHVN45) prior to LC-MS/MS acquisition.

### LC-MS/MS data acquisition and data processing

Peptides were analyzed using an Orbitrap Exploris 480 mass spectrometer (Thermo Fisher Scientific) coupled to a Vanquish Neo UHPLC system (Thermo Fisher Scientific). Peptide separation was performed on a PepSep reversed-phase C18 column (1.5 μm particles, 15 cm × 150 μm ID; Bruker) using a binary solvent system consisting of mobile phase A (0.1% formic acid in water) and mobile phase B (80% acetonitrile, 0.1% formic acid). The gradient was as follows: 3-28% B over 87 min, 28-40% B over 5 min, followed by a wash at 95% B for 8 min, at a flow rate of 600 nL/min. Data were acquired using data-dependent acquisition (DDA). Full MS scans (MS1) were acquired in the Orbitrap at a resolution of 120,000 (at *m/z* 200) over a mass range of 350-1,250 *m/z*, with a normalized AGC target of 400% and a dynamic exclusion of 45 s. The MS2 cycle time was set to 1 s. Precursors were isolated using a 1.6 *m/z* isolation window and fragmented by higher-energy collisional dissociation (HCD) at a normalized collision energy of 28%. MS2 spectra were acquired at a resolution of 15,000 using an AGC target of 300% in Auto Scan Range mode. Raw data were processed using FragPipe (version 23.1)^10^. Spectra were searched against the UniProt human canonical protein database (downloaded February 23, 2026; 20,416 entries). Search parameters included trypsin specificity with up to two missed cleavages. Carbamidomethylation of cysteine was set as a fixed modification, while methionine oxidation, protein N-terminal acetylation, and phosphorylation of serine, threonine, and tyrosine were set as variable modifications. IonQuant (version 1.11.20) was used to quantify phosphopeptides across all runs^11^. The PTMprophet (version 7.3.0) site localization threshold was set to 0.75^12^. A minimum peptide length of seven amino acids was required. Peptide and protein identifications were filtered at a 1% false discovery rate (FDR).

### Phosphosite quantification and differential analysis

Phosphopeptide intensities were processed with artMS 1.29.0 and MSstats 4.20.0 in R 4.6.1. Contaminants were removed and the analysis was restricted to phosphorylated (STY) features. The quantitative unit was the phosphosite feature, defined as a UniProt accession together with the modified residue and its position; phosphopeptides carrying more than one phosphosite were retained as composite features throughout, and were used only where unambiguous single-site assignment was not required. Intensities were log2-transformed, normalized across runs by median equalization and summarized per feature by Tukey’s median polish, with all features retained and no removal of outlier or uninformative features. No imputation was performed at any stage and censored intensities were treated as missing. This yielded 26,127 phosphosite features across the 27 runs, of which 4,463 were quantified in every sample.

Two families of contrasts were tested. The response to RMC-6236 was tested within each clone as RMC-6236 versus DMSO both at 4h and at 24h. Resistance was also tested as difference-in-differences interaction contrasts as linear combinations of the condition means in the MSstats contrast matrix that compares each edited clone with the unedited clone and the two edited clones with one another, at both time points (for example (H1047R RMC 4 h - H1047R DMSO) - (unedited RMC 4 h - unedited DMSO). Interaction log2 fold changes are hence differences between treatment responses and not differences in phosphosite abundance between clones.

### Kinase activity inference

Kinase and regulator activities were inferred from the single-clone responses by kinase–substrate enrichment analysis (KSEA)^13^. Kinase-substrate relationships were taken from the curated PhosphoSitePlus entries of the KSData table of the KSEAapp R package, version 2.0 ^14^. The same table also carries 10,000 relationships predicted by NetworKIN, which were not used. Of the curated entries, 21 were discarded for lacking either a serine, threonine or tyrosine position or a substrate gene symbol, leaving 9,736 relationships covering 348 kinases. Scores were computed from the MSstats log2 fold changes rather than with the package’s own implementation. Within each contrast, substrate sets and the background against which they were compared were drawn from the same set of single site phosphosites, and regulators with fewer than five quantified substrates were not tested. Two-sided P values from the standard normal distribution were corrected by Benjamini–Hochberg within each contrast. Of the 47 regulators tested, the 19 reaching an adjusted P value below 0.05 in at least one of the six responses are shown in **Fig. 3C**.

### Targeted mTORC1 substrate panel

The phosphorylation of 4E-BP1 was read directly from the single-clone MSstats contrasts. Sites shown in **Fig. 3D** are the canonical mTORC1 phosphorylation sites of 4E-BP1^15^ together with any other quantified 4E-BP1 phosphosite reaching an adjusted P value of 0.10 or below in at least one contrast.

### Pathway over-representation analysis

Proteins carrying significantly regulated phosphosites in the interaction contrasts were collapsed to gene symbols, so that a protein contributes once irrespective of the number of regulated sites, and the lists of increased and of decreased differential responses were submitted separately to Enrichr. Over-representation against the BioPlanet 2019 ^16^ library was assessed by one-sided Fisher exact tests against the Enrichr background of 20,000 annotated genes, with Benjamini-Hochberg correction within each gene list. Pathways were grouped into categories by the genes carrying their overlap rather than by name.

### Statistics and reproducibility

Three independent biological replicates were prepared per condition, and after QC no samples or data points were excluded. P values from the MSstats model-based tests were adjusted within each contrast by the Benjamini–Hochberg procedure, and phosphosites were considered significantly regulated at an adjusted P value below 0.05 together with |log2 fold change| ≥ log2(1.5). These two criteria define significance in all phosphosite-level panels of Fig. 3 and of the supplementary figures related to figure 3.

## RESULTS

### *PIK3CA* MUTATION CAUSES RESISTANCE TO KRAS INHIBITORS IN PATIENT-DERIVED CRC CELL LINES

Analysis of clinical sequencing data from a colon adenocarcinoma cohort (TCGA, GDC 2025, n = 429 sequenced samples) showed that *KRAS* is mutated in 42.3% of tumors, and that about 40% of *KRAS*-mutant tumors carry a concurrent *PIK3CA* mutation (**Fig. 1A**). *PIK3CA* mutations in these co-mutated tumors predominantly appear in the helical domain, at E545K (26.8%) and E542K (12.7%), or at the kinase-domain hotspot H1047R (9.9%) (**Fig. 1A**). Because KRAS and PI3K signaling converge on overlapping effectors, the MAPK cascade downstream of KRAS and the AKT/mTORC1 axis downstream of PI3K both ultimately drive proliferative and survival programs (**Fig. 1B**). Hence, we hypothesized that *PIK3CA* co-mutation might provide tumor cells with an escape route from KRAS-directed therapy.

**Figure 1:**
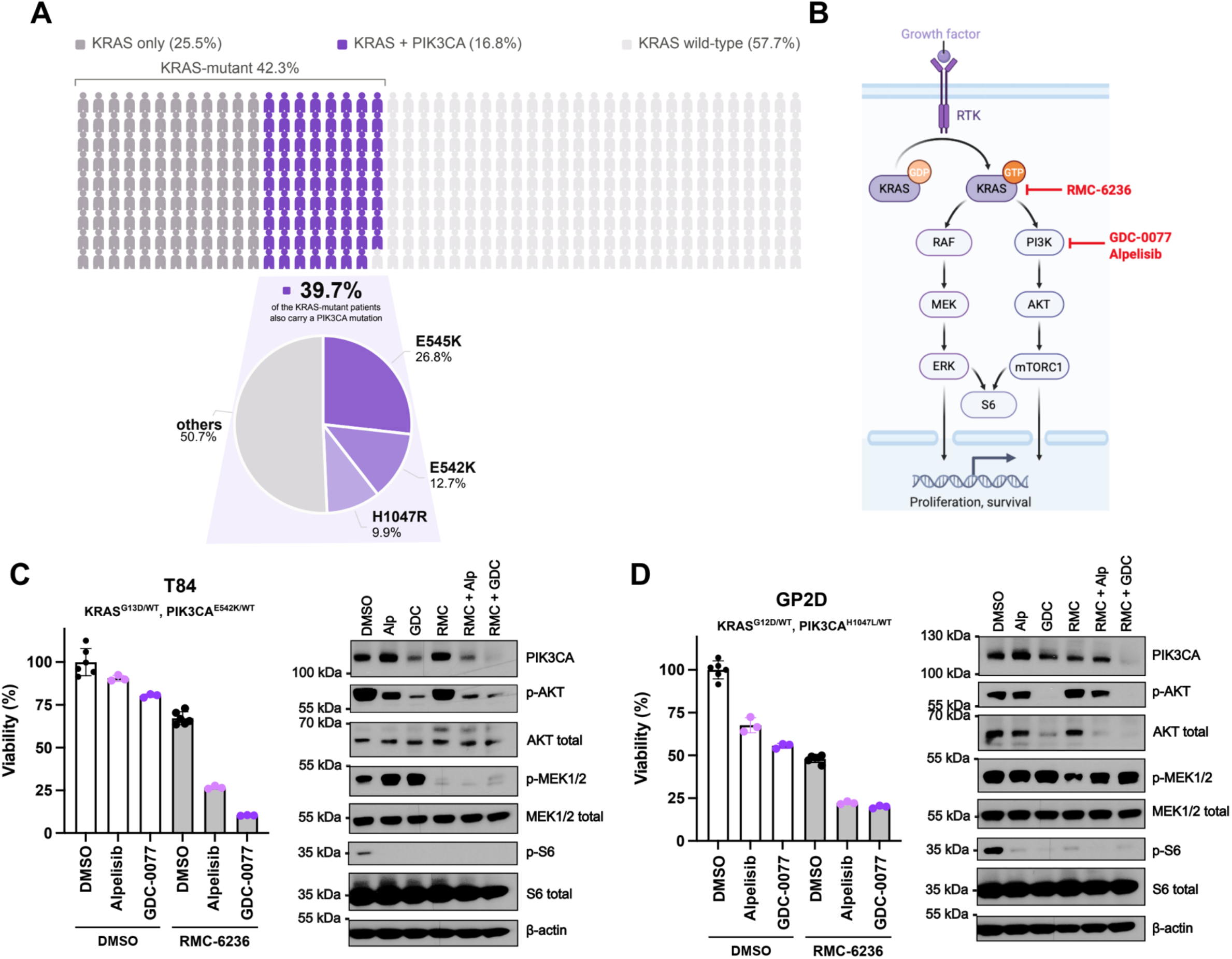
PIK3CA mutation causes resistance to KRAS inhibitors in patient-derived CRC cell lines. (**A**) Prevalence of *KRAS* and *PIK3CA* mutations in a colon adenocarcinoma cohort (TCGA, GDC 2025 release; cBioPortal study ID coad_tcga_gdc, n = 429 sequenced samples). *KRAS* is mutated in 42.3% of tumors, of which around 40% also carry a *PIK3CA* mutation. Pie chart: distribution of PIK3CA mutations among *KRAS*/*PIK3CA* co-mutated tumors: E545K (26.8%), E542K (12.7%), H1047R (9.9%), and other *PIK3CA* alterations (50.7%). (**B**) Schematic of the KRAS/MAPK and PI3K/AKT/mTORC1 signaling axes downstream of receptor tyrosine kinase (RTK) activation, showing the respective inhibitory targets of the pan-RAS(ON) inhibitor Daraxonrasib (RMC-6236) and the PI3Kα inhibitors inavolisib (GDC-0077) and alpelisib, created with BioRender.com. (**C-D**) *KRAS*/*PIK3CA* co-mutated T84 (*KRAS^G13D/WT^*, *PIK3CA^E542K/WT^) (***C**) and GP2D (*KRAS^G12D/WT^*, *PIK3CA^H1047L/WT^*) (**D**) CRC cells were treated for 24h with alpelisib (500 nM) or GDC-0077 (500 nM), alone or in combination with RMC-6236 (500 nM). Left, viability by CellTiter-Glo normalized to DMSO; bars show mean ± SD (n = 3-6 replicates), individual values overlaid (n = 1 representative experiment). Right, immunoblot of PIK3CA, phospho- and total AKT, phospho- and total MEK1/2, phospho- and total S6, and β-actin (loading control) (n = 2 independent experiments).

To test this, we treated two *KRAS/PIK3CA* co-mutated CRC cell lines, T84 (*KRAS^G13D/WT^*, *PIK3CA^E542K/WT^*) and GP2D (*KRAS^G12D/WT^*, *PIK3CA^H1047L/WT^*), with the pan-RAS(ON) inhibitor RMC-6236, alone or in combination with the PI3Kα inhibitors alpelisib or inavolisib (GDC-0077), the latter also acting as a degrader of mutant PIK3CA^17^ (**Fig. 1C-D**). Doses for each agent were selected based on prior dose-response curves in these cell lines, using the IC25 of each compound. RMC-6236 treatment alone effectively shut down KRAS/MAPK output, reducing phospho-MEK1/2 in both lines, but had only a modest effect on viability. Notably, GDC-0077 more effectively suppressed phospho-AKT and phospho-S6 as a single agent than alpelisib, particularly in GP2D, consistent with its additional activity as a degrader of mutant PIK3CA. More importantly, combining RMC-6236 with alpelisib or GDC-0077 abolished residual phospho-AKT and phospho-S6 signaling and drove a synergistic loss of viability in both cell lines, significantly exceeding either PI3Kα inhibitor alone (**Fig. 1C-D**). Together, these data indicate that *PIK3CA* co-mutation sustains AKT/mTORC1 signaling and cell viability under KRAS inhibition, and that dual KRAS/PI3Kα blockade is required to achieve effective growth suppression in *KRAS*/*PIK3CA* co-mutated CRC.

### *PIK3CA*/*KRAS* CO-MUTATION CAUSES RESISTANCE TO KRAS INHIBITION IN ISOGENIC CRC MODELS

To generate genetically defined models of KRAS/PIK3CA co-mutation, we engineered isogenic *KRAS*/*PIK3CA* co-mutant CRC cell lines using two complementary approaches: doxycycline-inducible over-expression of FLAG-tagged PIK3CA (WT, H1047R, or E545K), and CRISPR-Cas9 edition of the endogenous *PIK3CA* hotspot mutations in KRAS-mutant CRC lines selected to span the KRAS mutational landscape of CRC. KRAS G12D and G12V are the two most frequent KRAS alleles in CRC, which together account for approximately half of KRAS-mutant tumors. Although KRAS G12C is less prevalent, it represents the first KRAS allele for which selective covalent inhibitors have reached clinical approval (sotorasib, adagrasib)^18^. We therefore selected SW837 (KRAS^G12C/WT^) and SW620 (KRAS^G12V/WT^) to generate isogenic models which cover both the most prevalent and the most clinically actionable KRAS genotypes in CRC (**Fig. 2A**).

**Figure 2:**
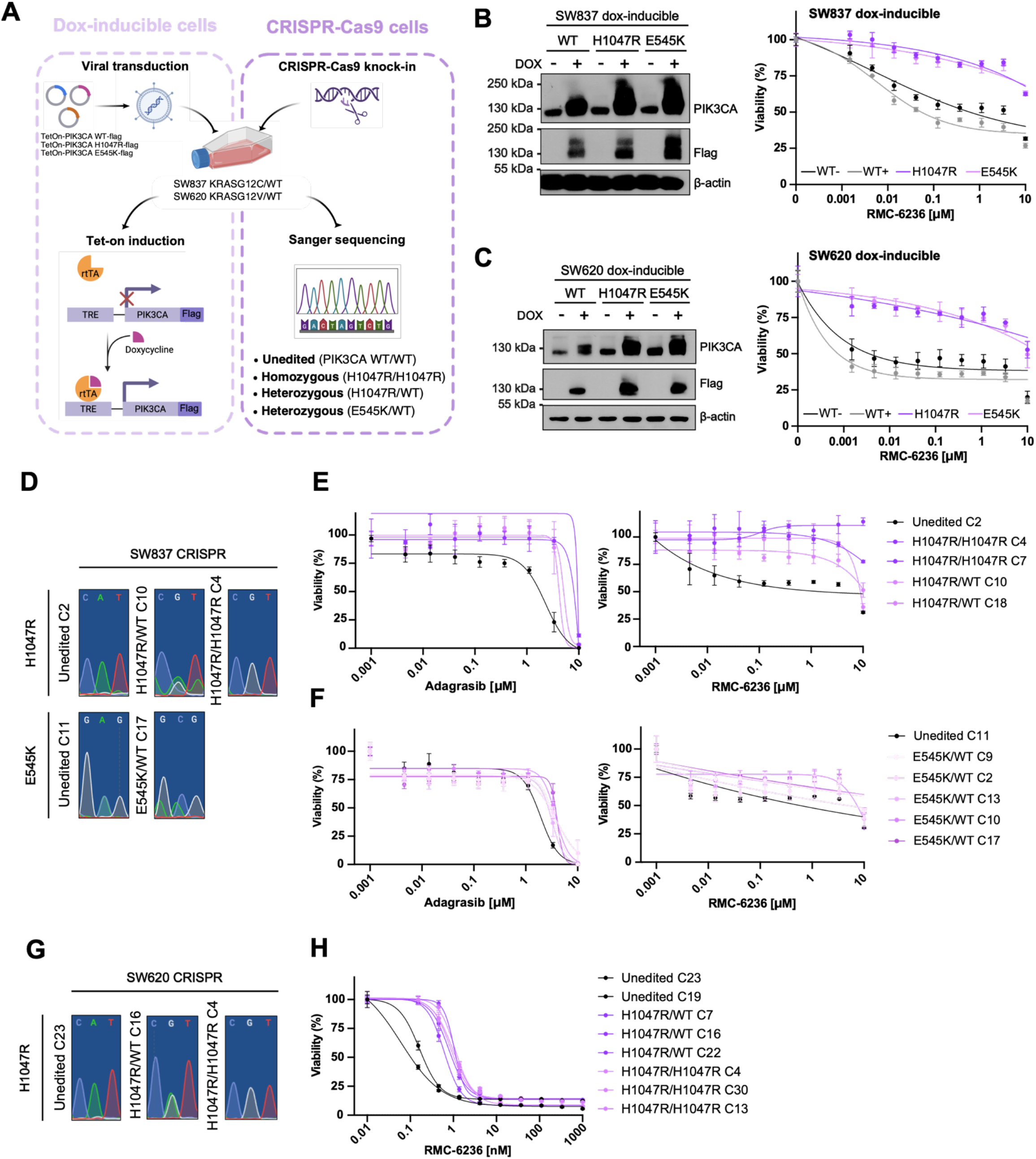
PIK3CA/KRAS co-mutations cause resistance to KRAS inhibitor in isogenic CRC models. **A.** Schematic representation of the generation of isogenic *KRAS/PIK3CA* co-mutant CRC model generation. Left: doxycycline-inducible FLAG-tagged PIK3CA WT, H1047R, or E545K constructs were stably introduced by viral transduction. Right: PIK3CA H1047R or E545K edited by CRISPR-Cas9, validated by Sanger sequencing. Both approaches were applied to SW837 (*KRAS G12C*) and SW620 (*KRAS G12V)* cell lines**. B-C.** SW837 (B) and SW620 (C) Dox-inducible FLAG-PIK3CA cells were treated with doxycycline to induce WT, H1047R, or E545K PIK3CA-FLAG expression (immunoblot, left), were treated with increasing doses of RMC-6236 for 5 days and subjected to CellTiter-Glo viability assay (right). Data are mean ± SD of n = 3 independent experiments. WT−, WT+: uninduced and doxycycline-induced WT PIK3CA, H1047R PIK3CA dox-induced or E545K PIK3CA dox-induced. **D.** Sanger sequencing chromatograms confirming unedited (WT/WT), homozygous, and heterozygous CRISPR-Cas9 PIK3CA H1047R or E545K edited clones in SW837 cells**. E-F.** SW837 PIK3CA H1047R (E) or E545K (F) CRISPR edited clones were treated with increasing doses of adagrasib (left) or RMC-6236 (right) for 5 days and viability was assessed by CellTiter-Glo (n = 2 independent experiments). **G.** Sanger sequencing chromatograms confirming unedited and heterozygous/homozygous PIK3CA H1047R CRISPR-Cas9 edited clones in SW620 cells. **H.** SW620 unedited and PIK3CA H1047R (heterozygous and homozygous) clones were treated with increasing doses of RMC-6236 for 5 days and viability was assessed by CellTiter-Glo (n = 2 independent experiments).

**Figure 3:**
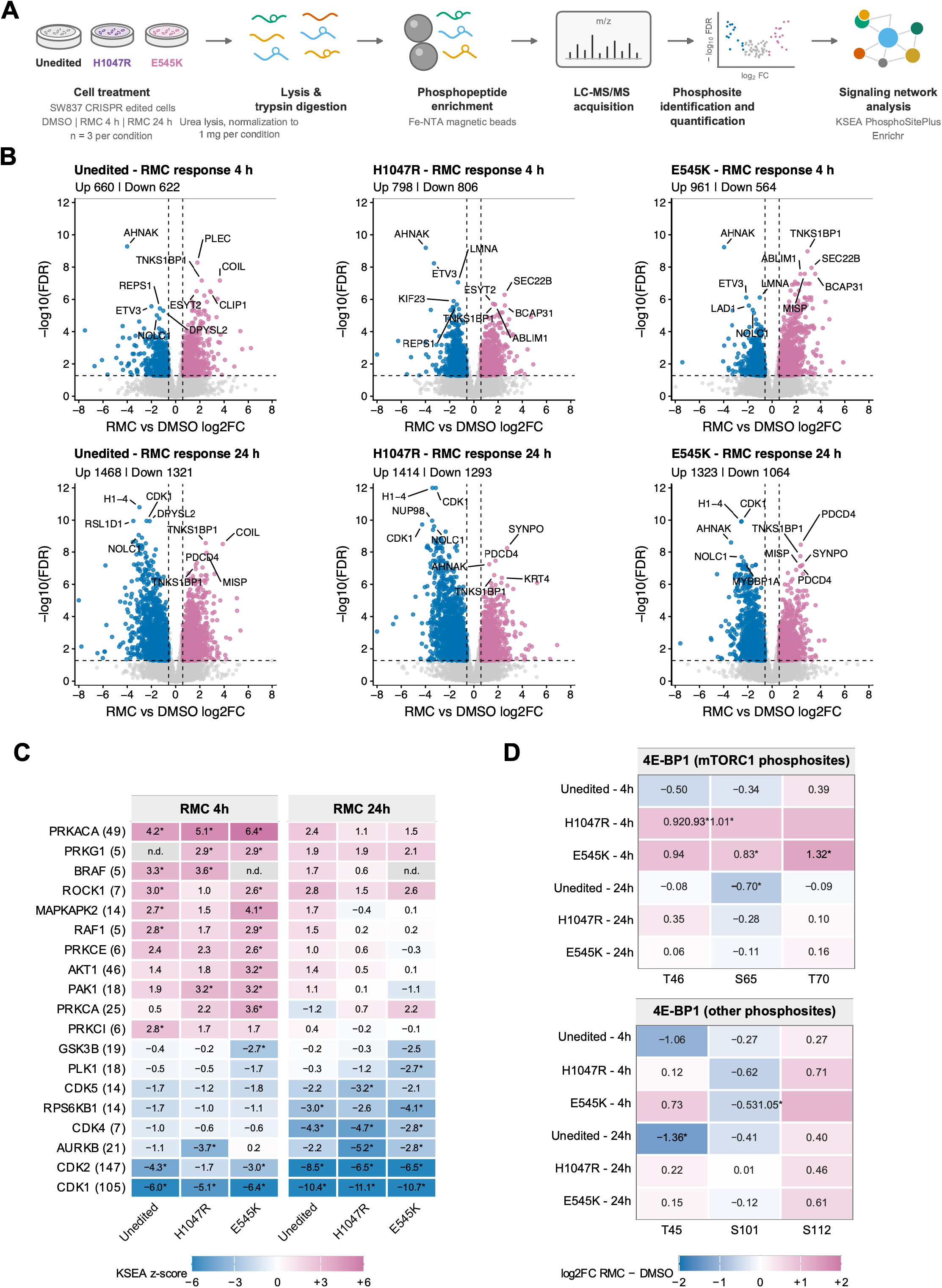
Phosphoproteomic response to RAS inhibition in *PIK3CA* WT and mutant CRC cell lines. **A.** Global phosphoproteomic profiling workflow. SW837 CRISPR clones (unedited clone 2 and CRISPR-Cas9 edition of PIK3CA H1047R clone 10 or E545K clone 17) were treated with RMC-6236 (100 nM) or DMSO for 4 or 24 h. Phosphopeptides were enriched on Fe-NTA IMAC beads and analyzed by LC-MS/MS. n = 3 biological replicates per condition (27 runs total). **B.** Volcano plots of the six single-clone responses to RMC-6236 (RMC-6236 vs. DMSO within each clone, at 4 and 24 h). x: log2 fold change; y: −log10(FDR); dashed lines: FDR = 0.05 and |log2FC| = log2(1.5); pink: increased, blue: decreased, grey: not significant. Numbers of significant sites are given on top of each panel. The five most significant increases and decreases of each contrast are labelled by gene symbol, ranked by adjusted P value. **C.** Kinase-substrate enrichment analysis of the same six responses, using PhosphoSitePlus kinase-substrate relationships. Color and values: KSEA z-score (pink: increased inferred activity; blue: decreased inferred activity; color scale capped at ±6). Asterisk: inferred activity significantly different from DMSO within that clone (FDR < 0.05, Benjamini–Hochberg within each contrast). The number in brackets after each regulator is m (the number of annotated substrates quantified), given as the median over the six contrasts. n.d.: fewer than five substrates quantified, so the regulator was not tested in that contrast. Regulators are ordered by mean z and are all of those reaching FDR < 0.05 in at least one of the six contrasts. **D.** RMC-6236 response of EIF4EBP1 (4E-BP1) phosphosites in each of the six single-clone contrasts. Values and fill are MSstats log2 fold changes of RMC-6236 versus DMSO within each clone and time point (pink: increased, blue: decreased), scale capped at ±2. Upper block, canonical mTORC1 sites of 4E-BP1, lower block: other quantified 4E-BP1 phosphosites reaching FDR ≤ 0.10 in at least one contrast. Asterisk: FDR < 0.05 and |log2FC| ≥ log2(1.5).

In the dox-inducible SW837 and SW620 models, expression of WT PIK3CA modestly decreased sensitivity to the pan-RAS(ON) inhibitor RMC-6236 relative to the uninduced condition, whereas induction of either PIK3CA H1047R or E545K conferred a marked loss of sensitivity to RMC-6236 in both cell lines (**Fig. 2B-C**). This was recapitulated in the CRISPR-Cas9 edited models: SW837 clones homozygous or heterozygous for PIK3CA H1047R were markedly less sensitive than their isogenic unedited counterparts to both adagrasib and RMC-6236 (**Fig. 2D-E**). Interestingly, heterozygous PIK3CA E545K clones showed a qualitatively similar, but more modest, loss of sensitivity to both inhibitors (**Fig. 2F**). The resistance phenotype conferred by PIK3CA H1047R was independently reproduced in SW620 edited clones (heterozygous and homozygous), which both remained markedly less sensitive to RMC-6236 than unedited controls (**Fig. 2G-H**). Together, these isogenic models demonstrate that in matched genetic backgrounds, PIK3CA hotspot mutations are sufficient to drive resistance to KRAS inhibition in co-mutated CRC cells.

### ONCOGENIC PIK3CA RESHAPES THE PHOSPHOPROTEOMIC RESPONSE TO RAS INHIBITION

To better define the signaling basis of resistance in *KRAS*/*PIK3CA* co-mutated cells, we profiled the global phosphoproteome of unedited and CRISPR-edited PIK3CA H1047R and PIK3CA E545K SW837 clones after 4 h or 24 h of RMC-6236 (100 nM) or DMSO, in three biological replicates per condition (**Fig. 3A**). Sample-to-sample correlation was uniform across runs, and phosphopeptides accounted for 78.9–97.2% of identified peptides (median: 95.1%); only three runs fell below 90%, and all were retained for the analysis. The experiment quantified 26,127 phosphosite features, of which 4,463 were measured in every one of the 27 runs. (**Supp. data 1A-B**).

Of the shared quantified phosphosites between comparisons, 1,282 to 1,604 phosphosites per clone responded significantly to treatment at 4 h, and 2,387 to 2,789 phosphosites at 24 h (with FDR < 0.05 and fold change ≥ 1.5, log2FC ≥ 0.58). The five most significant hits that increased and decreased from each contrast are labelled on each respective volcano plot (**Fig. 3B**).

To identify plausible genotype-dependent differential regulation of kinases, we ran a Kinase-substrate enrichment analysis (KSEA) using the PhosphositePlus resource^13, 14^. KSEA analysis of the six single-clone responses recovered the expected pharmacology of the drug: kinases involved in cell cycle progression such as CDK and aurora kinase activity fell in every clone, most strongly at 24 h, while PKCα (PRKCA), RAF and ROCK signaling increased at 4 h (**Fig. 3C**). For most kinases and other regulators inferred by KSEA the activity profile was the same in the unedited and in the mutant clones. Indeed, in 64 of the 72 comparisons (DMSO versus RMC-6236) that could be made, the drug shifted inferred activity in the same direction, with PIK3CA mutation only partially attenuating or augmenting the response. Still, three kinases were inferred to behave differently: AKT1 activation reached significance in the E545K clone at 4 h and at no other clone or time point, as did PRKCA, while GSK3B, which AKT1 inhibits, fell at the same point. Corroborating these results, we see that at 4 h the mTORC1 target sites S65 and T70 of 4E-BP1 gained phosphorylation in both mutant clones (S65: +0.93 and +0.83 log2FC, T70: +1.01 and +1.32 log2FC) while remaining unchanged in the unedited clone (−0.34 and +0.39 log2FC; **Fig. 3D**).

Principal component analysis (PCA) of the quantified phosphosites separated the samples along two independent axes (**Supp. Data 1C**). The first axis (25.8% of the variance) arranged the samples by treatment (vehicle, then 4 h, then 24 h) and in the same order in the three clones. The second axis (14.4%) separated the three clones from one another, even before treatment. The drug therefore moves every clone, but the clones do not start from the same phosphorylation state, hence supporting a comparison of RMC-6236 response as a difference of differences, rather than by comparing treated clones directly.

Hence, to identify differences in phosphorylation that might not be evident at the level of kinase inference, a pairwise interaction analysis of the RMC-6236 response was generated (mutant vs unedited, or E545K vs H1047R). This analysis compared the RMC-6236 response (RMC-6236 vs. DMSO) between genotypes: H1047R vs. unedited and E545K vs. unedited, and directly between the two mutant alleles (E545K vs. H1047R), separately at 4 h and 24 h. This evaluation revealed markedly asymmetric numbers of differentially responsive phosphosites across genotypes and time points (**Fig. 4A-B**). First, both H1047R and E545K PIK3CA mutant clones diverged from the unedited clone at both time points (H1047R, 120 sites at 4 h and 213 at 24 h; E545K, 100 sites at 4h and 129 sites at 24h), whereas the direct comparison between the two mutant clones yielded only 7 and 14 sites with no coherent functional signature (**Fig. 4A-B**). Despite their distinct genotypes, the two PIK3CA hotspot alleles therefore drove largely convergent resistance programs, while both differed substantially from the RMC-sensitive parental clone.

**Figure 4:**
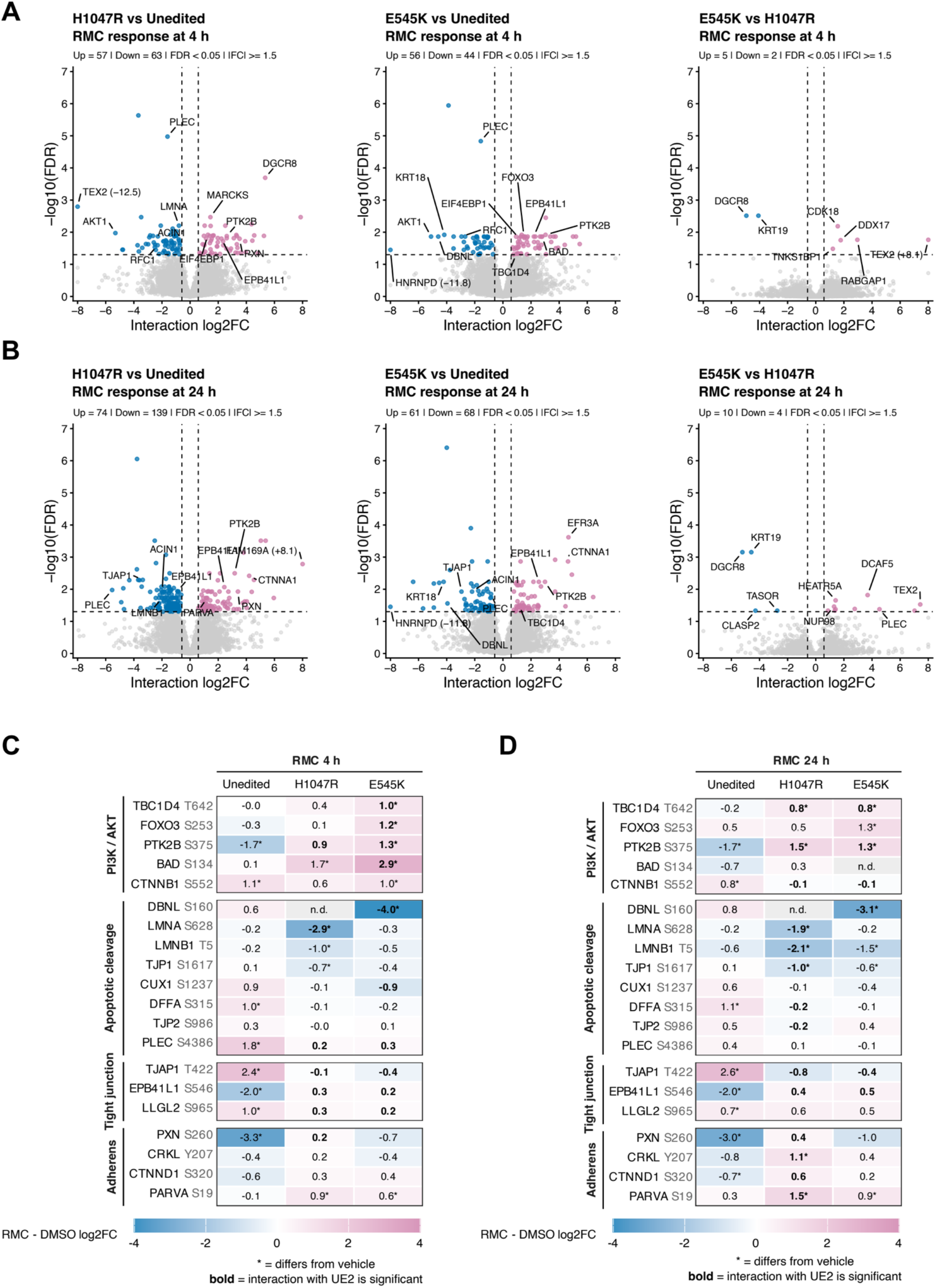
Oncogenic *PIK3CA* reshapes the phosphoproteomic response to RAS inhibition. A-B. Volcano plots of the three interaction contrasts (H1047R vs. unedited, E545K vs. unedited, and E545K vs. H1047R), at 4 (**A**) and 24 h (**B**). Interaction log2 fold changes compare treatment responses; positive values indicate a response more positive, or less negative, in the first-named clone than in the second. FDR < 0.05 and |log2FC| ≥ log2(1.5); pink: increased, blue: decreased, grey: not significant. The five most significant increases and decreases of each contrast are labelled by gene symbol, ranked by adjusted P value. **C-D.** RMC-6236 response of the phosphosites driving the interaction signal in the pathway over-represented among proteins carrying differentially responsive phosphosites in the mutant-vs-unedited contrasts of B (H1047R vs. unedited and E545K vs. unedited, each at 4 and 24 h), at 4 h (**C**) and 24 h (**D**), from gene-level lists tested with Enrichr (BioPlanet 2019 ^16^, default 20,000-gene background, Benjamini-Hochberg correction within each list). Printed values are RMC-vs-DMSO log2 fold change within each clone (scale capped at ±4; grey: not quantified). Asterisk: significant RMC-vs-DMSO response within that clone (FDR < 0.05, |log2FC| ≥ log2(1.5)). Bold: significant interaction (a response significantly different from the unedited clone) at the same thresholds.

We then performed a functional enrichment assessment for pathway over-representation using Enrichr. Pathway over-representation (Enrichr-BioPlanet ^16^) identified PI3K/AKT terms specifically in E545K at 4 h (under “nicotinic acetylcholine receptor role in the regulation of apoptosis” with FDR= 0.0034; “PIK3C1/AKT pathway” with FDR= 0.0095 and “PI3K/PLC/TRK pathway” with FDR= 0.0095 terms; **Fig. 4C-D**). At the site level, we also observed in the unedited clone a loss of PI3K/AKT substrate phosphorylation. Indeed, PTK2B S375 fell by 1.71 log2FC at 4 h and by 1.67 at 24 h in the unedited clone, whereas the same site rose in both mutant clones, by 0.90 and 1.32 at 4 h and by 1.50 and 1.32 at 24 h in H1047R and E545K clones respectively (**Fig. 4C-D**). Two AKT substrates behaved in the same way: TBC1D4 T642 increased significantly in E545K at 4 h (+0.96 log2FC) and in both mutants at 24 h (+0.76 and +0.76 log2FC). Similarly, FOXO3 S253 increased in E545K at both time points (+1.18 and +1.31 log2FC) and neither site moved in the unedited clone.

RMC-6236 therefore fails to switch off the PI3K-AKT-mTORC1 axis in either mutant clone, and the remaining differences between the clones can be read as the consequences of that failure at two distinct AKT effector arms.

First, pathway over-representation identified apoptotic cleavage terms in the mutant clones specifically at 24 h (under “Apoptosis execution phase” with FDR<0.0001 in H1047R clone and FDR=0.0047 in E545K clone; “Apoptosis” with FDR= 0.0037 in H1047R clone and “Apoptotic cleavage of adhesion proteins” with FDR= 0.0049 in H1047R clone terms; **Fig. 4C-D**). Moreover, at the site level, the phosphorylation of the BH3-only protein BAD at S134 increased by 1.70 log2FC in H1047R and by 2.94 in E545K at 4 h, while the unedited clone did not respond (+0.14 log2FC; **Fig. 4C**). Phosphorylated BAD is sequestered by 14-3-3 proteins and can no longer neutralize BCL-2 and BCL-XL ^19, 20^ , so this represents an active suppression of the apoptotic machinery rather than an absence of response. FOXO3 S253 is the transcriptional counterpart of the same output, since AKT-mediated phosphorylation of this site excludes FOXO3 from the nucleus and silences its pro-apoptotic target genes ^21^. Both mutant clones thus engage two independent anti-apoptotic AKT outputs within 4 h of exposure.

Second, cell-adhesion signaling was remodeled in opposite directions at adherens/focal adhesions versus tight junctions. Indeed, pathway over-representation reported increased phosphorylation of adherens adhesion components in the mutant clones, especially in the H1047R clone (under “Cell-cell communication” with FDR=0.0067; “Cell junction organization” with FDR= 0.0085 and Cell-extracellular matrix interactions with FDR=0.0213; **Fig. 4C-D**). At the site level, the adhesion kinase PTK2B is the AKT substrate with the largest divergence between genotypes (-1.7 log2FC in unedited versus +0.9 and 1.3 log2FC in H1047R and E545K). Additionally, phosphorylation of PARVA S19, a component of the ILK-PINCH-parvin complex of focal adhesions, increased in both mutant clones at both time points (+0.87 and +0.59 log2FC at 4 h, +1.48 and +0.89 at 24 h) while the unedited clone did not respond, and CRKL Y207 increased in H1047R at 24 h (+1.07 log2FC). In addition, the dephosphorylation of PXN S260 (−3.31 and −3.02 log2FC) and CTNND1 S320 (−0.69 log2FC) that RMC-6236 induced in the unedited clone was absent in both mutants (**Fig. 4C-D**). Phosphorylation of tight junction components moved in the opposite direction: this term was instead enriched among sites whose response was more negative in the mutant clones (under “Tight junction” with FDR=0.0173 in E545K clone at 4 h and FDR=0.0139 in H1047R clone at 24 h) (**Fig. 4C-D**). Indeed, the unedited clone remodeled these “tight junction” sites in both directions under RMC-6236, gaining phosphorylation on TJAP1 T422 (+2.4 and +2.6 log2FC) and LLGL2 S965 (+1.0 and +0.7 log2FC), and losing it on EPB41L1 S546 (−2.0 and −2.0 log2FC), while neither mutant clone showed either change. The resistant clones are thus distinguished by an absence of tight-junction remodeling, alongside a gain of adherens-junction/focal-adhesion signaling.

Together, these data support a time-resolved model in which PIK3CA hotspot mutations preserve AKT substrate phosphorylation under RAS inhibition, suppressing the pro-apoptotic response to RMC-6236. E545K engages this program earliest, with its PI3K/AKT signature and largest BAD S134 response converging at 4 h, while H1047R shows a delayed but broader remodeling that peaks at 24 h and extends to junction and adhesion phosphosites. This difference in timing means the 4 h data capture an early signaling effect, whereas the 24 h data may also reflect the resulting divergence in cell fate.

### INHIBITION OF PIK3CA POTENTIATES RMC-6236 EFFECTS IN *PIK3CA/KRAS* CO-MUTATED CRC

We next asked whether dual KRAS/PI3Kα inhibition could rescue *PIK3CA* mutation-driven resistance to RMC-6236, building on our observation in **Fig. 1C-D** that combining either PI3Kα inhibitor (alpelisib or GDC-0077) with RMC-6236 significantly reduced viability in two co-mutated CRC cell lines. In our isogenic models, and consistent with **Fig. 2B-C**, doxycycline-induced overexpression of WT PIK3CA caused only a slight decrease in sensitivity to RMC-6236 relative to the no-dox control, whereas overexpression of either PIK3CA H1047R or E545K caused a large, significant decrease in sensitivity in both SW837 and SW620 dox-inducible cells (**Supp. Fig. 5A and Fig. 5A**). However, the combination of RMC-6236 with alpelisib did not rescue this resistance, while GDC-0077 did (**Fig. 5A**). We therefore chose GDC-0077 for subsequent experiments, hypothesizing that this difference reflects GDC-0077’s additional activity as a degrader of mutant PIK3CA, which alpelisib lacks. Similarly, in the SW837 and SW620 CRISPR Cas9 edited cell lines, the RMC-6236/GDC-0077 combination produced a substantial loss of viability (**Supp. Fig. 5B-C**). The strength of the synergy between RMC-6236 and GDC-0077 was quantified using a Bliss score (synergy if > 10%, additivity between −10% and 10%, antagonism if < −10%, following ^22^). In the SW837 CRISPR cell line, Bliss scores reached up to 20% in H1047R and 25% in E545K clones (vs. a maximum of 14% in unedited cells, where only four combinations were just above the synergy threshold) (**Fig. 5B**). Similarly, and more strikingly, in the SW620 CRISPR cells, Bliss scores reached up to 28% in SW620 H1047R clones (vs a maximum of 5% in unedited cells) in **Fig. 5C**. Synergy was hence observed across a broad range of RMC-6236/GDC-0077 dose combinations specifically in the PIK3CA-mutated clones, indicating that the strong synergy between these two compounds is restricted to a PIK3CA-mutant genetic background.

**Figure 5:**
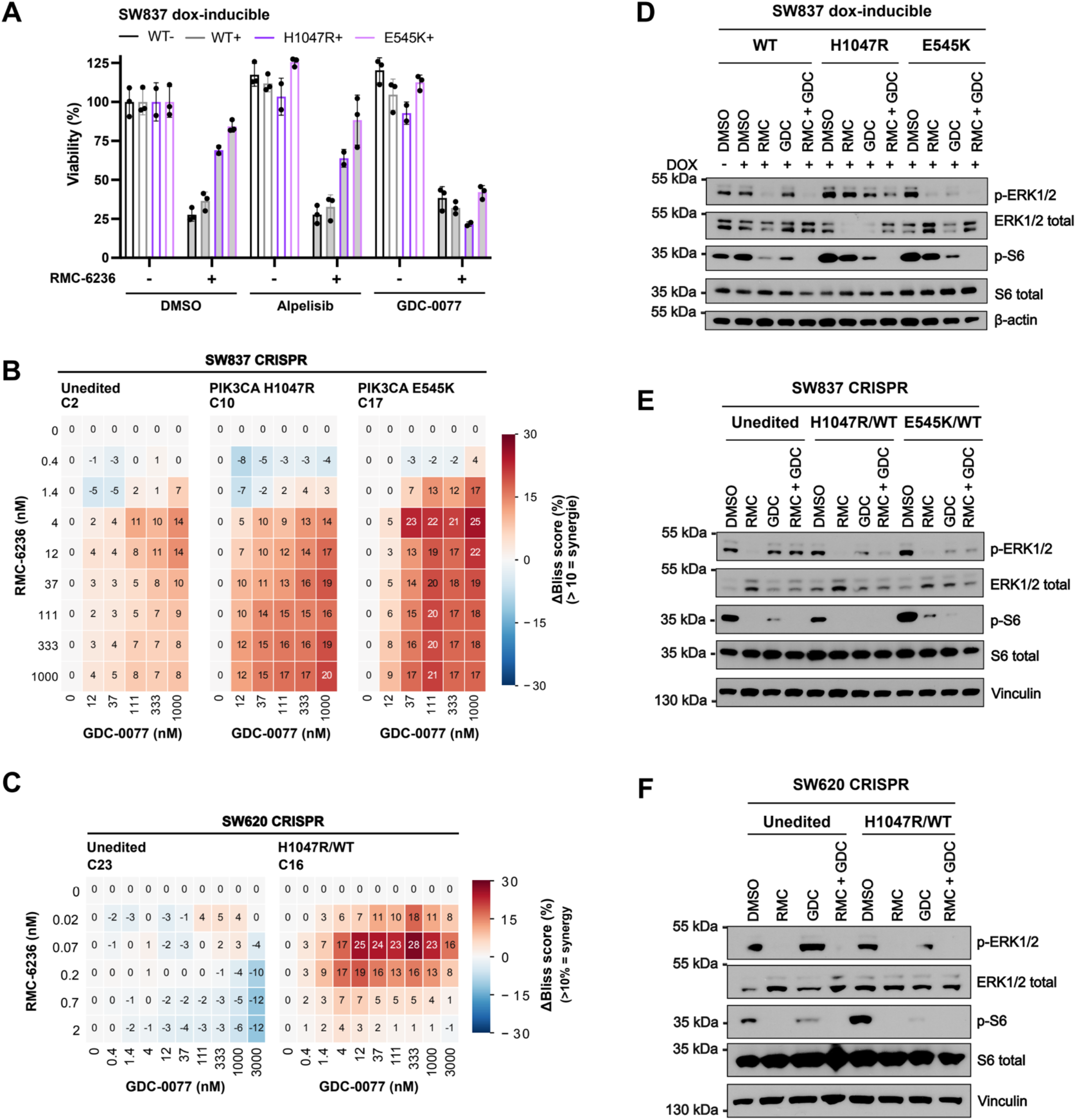
Inhibition of PIK3CA potentiates RMC-6236 effects in *PIK3CA/KRAS* co-mutated CRC. **A.** SW837 dox-inducible PIK3CA (WT, H1047R, or E545K)-expressing cells were treated with RMC-6236 (100 nM) with or without alpelisib (200 nM) or GDC-0077 (200 nM) for 5 days, cell viability was assessed by CellTiter-Glo. −/+: uninduced and doxycycline-induced PIK3CA. Data are mean ± SD of n = 2 independent experiments. **B**. SW837 CRISPR unedited (UE2), PIK3CA H1047R (clone 10), or PIK3CA E545K (clone 17) clones were treated with a dose matrix of RMC-6236 (0-1000 nM) and GDC-0077 (0-1000 nM) for 5 days and then subjected to CellTiter-Glo viability assay. Matrix-response synergy was quantified using a Bliss score (synergy if > 10%, additivity between 0–10%, antagonism if < 0%), see Supp. Fig. 5B for the corresponding viability matrixes. n = 3 independent experiments. **C**. SW620 CRISPR unedited (clone 23) or PIK3CA H1047R (clone 16) clones were treated with a dose matrix of RMC-6236 (0-1000 nM) and GDC-0077 (0-3000 nM) for 5 days and then subjected to CellTiter-Glo viability assay. Matrix-response synergy was quantified using a Bliss score (synergy if > 10%, additivity between 0–10%, antagonism if < 0%), see Supp. Fig. 5C for the corresponding viability matrixes. n = 2 independent experiments. **D-F**. Immunoblot of KRAS/MAPK and PI3K/AKT downstream signaling in response to RMC-6236 (100 nM) and GDC-0077 (200 nM), alone or in combination, for 24 h in SW837 dox-inducible (**D**), SW837 CRISPR (**E**), and SW620 CRISPR (**F**) cells. Loading controls: β-actin or vinculin.

At the signaling level, RMC-6236 (24 h) reduced phospho-ERK1/2 in SW837 dox-inducible (**Fig. 5D**), SW837 CRISPR (**Fig. 5E**), SW620 CRISPR (**Fig. 5F**) but also in SW620 dox-inducible (**Supp. Fig. 5D)** cells, confirming effective MAPK pathway inhibition, while GDC-0077 reduced phospho-S6 across the same models. Notably, neither RMC-6236 nor GDC-0077 alone reduced phospho-AKT in SW620 dox-inducible cells, but their combination did (**Supp. Fig. 5D**). This suggests that single-node inhibition of either KRAS or PI3K is insufficient to overcome compensatory feedback reactivation of AKT, and that durable AKT suppression and the resulting loss of viability may require blocking both nodes simultaneously.

Together, these data show that pharmacological inhibition of mutant *PIK3CA* restores sensitivity to RAS(ON) inhibition, coinciding with suppression of AKT phosphorylation that otherwise persists under either single agent alone. The RMC-6236/GDC-0077 combination thus emerges as a rationale strategy to overcome PIK3CA-driven resistance to KRAS inhibitors in colorectal cancer.

### RMC-6236 AND GDC-0077 COMBINATION TRIGGERS CRC CELL APOPTOSIS

Having shown that pharmacological PIK3CA inhibition restores RMC-6236 sensitivity, we next asked whether this combination predominantly drives cell death or instead induces a senescence-like cell growth arrest.

T84, GP2D, and dox-inducible SW620 and CRISPR SW620 cells were treated with RMC-6236 or GDC-0077, alone or in combination, and cell death was quantified using SYTOX-green live-cell imaging, where SYTOX-green selectively stains the nucleic acids of membrane-compromised, apoptotic cells. In T84 and GP2D cells, RMC-6236 and GDC-0077 monotherapy induced only a mild increase in cell death relative to DMSO (T84: RMC ∼15%, GDC ∼11%; GP2D: RMC ∼21%, GDC ∼33%), whereas the combination produced a substantially larger increase in both cell lines (T84: ∼42%; GP2D: ∼52%) (**Supp. Fig. 6A-B)**.

**Figure 6:**
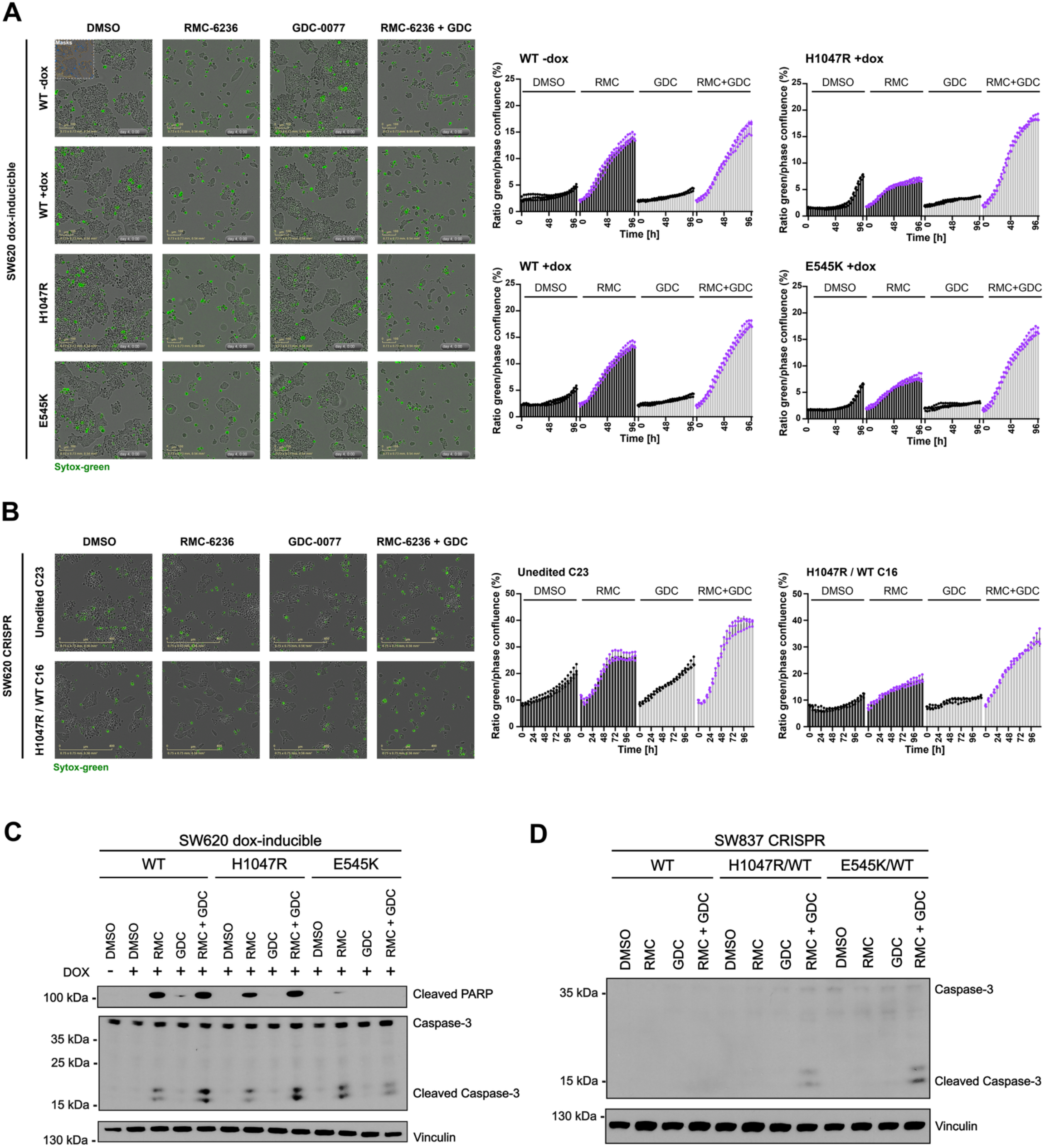
RMC-6236 and GDC-0077 combination triggers CRC cell apoptosis. **A.** SYTOX-green cell death detection by live-cell microscopy in SW620 dox-inducible cells (WT −dox, WT +dox, H1047R, E545K) treated with RMC-6236 (100 nM) with or without GDC-0077 (200 nM) for 5 days. Representative images of each condition at Day 5 with phase and green fluorescence combined are shown. The ratio of SYTOX-green-positive area to phase confluence is plotted over time for each condition (%). Scale bars: 100 μM. **B.** SYTOX-green cell death detection by live-cell microscopy in SW620 CRISPR cells (WT −dox, WT +dox, H1047R, E545K) treated with RMC-6236 (100 nM) with or without GDC-0077 (200 nM) for 5 days. Representative images of each condition at Day 5 with phase and green fluorescence combined are shown. The ratio of SYTOX-green-positive area to phase confluence is plotted over time for each condition (%). Scale bars: 400 μM. **C-D.** Immunoblot of the apoptosis markers cleaved PARP and cleaved caspase-3 in response to RMC-6236 (100 nM) and GDC-0077 (200 nM), alone or in combination, for 24 h in SW620 dox-inducible (**C**) or SW837 CRISPR (**D**) cells.

In dox-inducible SW620 cells (**Fig. 5A**), GDC-0077 alone did not increase cell death above DMSO baseline in any of the four genotypes (WT −dox, WT +dox, H1047R, E545K). RMC-6236 alone induced moderate cell death in cells with dox-inducible WT PIK3CA (∼14% at 96 h, both −dox and +dox) but a smaller increase in the PIK3CA-mutant clones (∼7%, roughly half of the WT response). The combination further increased cell death in the mutant clones (∼19%, approximately 2.5-fold over RMC-6236 alone), whereas in WT cells it produced only a modest (∼17%) additional increase over RMC-6236 alone. Similarly, in SW620 CRISPR cells, GDC-0077 alone did not increase cell death above DMSO baseline. However, consistent with our previous results, RMC-6236 alone induced moderate cell death in the unedited cells and no difference as compared to DMSO in the H1047R/WT clone. Notably, the combination of both GDC-0077 and RMC-6236 resulted in a mild increase in cell death in the unedited clone while it triggered almost a doubling of cell death induction as compared to RMC-6236 alone treatment in the H1047R/WT clone.

Apoptosis induction was confirmed by immunoblotting using cleaved PARP and cleaved caspase-3 as apoptosis markers. In T84 and GP2D cells, RMC-6236 or GDC-0077 alone caused only a modest increase in cleaved PARP and cleaved caspase-3, while the combination markedly increased both apoptosis markers (**Supp. Fig. 6C-D)**. In dox-inducible SW620 cells (**Fig. 6C**), the combination increased cleaved caspase-3 above RMC-6236 alone in WT and H1047R cells, but not in E545K cells. Cleaved PARP showed a more restricted pattern: it increased over RMC-6236 alone only in H1047R cells. Indeed, in WT cells, RMC-6236 alone already produced a strong signal that the combination did not further increase, and in E545K cells cleaved PARP was undetectable in the combination despite a faint signal with RMC-6236 alone (**Fig. 6C**). In SW837 CRISPR clones (**Fig. 6D**), the combination increased cleaved caspase-3 specifically in the PIK3CA-mutated clones (H1047R, E545K), with no detectable increase in the unedited clone.

Overall, these data indicate that the RMC-6236/GDC-0077 combination induces apoptosis preferentially in PIK3CA co-mutated CRC cells across multiple models.

## Discussion

Despite dramatic progress towards targeting KRAS with allele-specific and now pan-KRAS directed therapeutics, patients with CRC currently only have limited access to these agents. The reduced benefit seen with adagrasib for CRC patients with *KRAS(G12C)* mutation highlights the importance of understanding mechanisms of resistance for KRAS inhibition in this disease. Frequent *KRAS/PIK3CA* co-mutation is an important potential mechanism of resistance for which understanding remains inadequate.

This study demonstrates that adding an oncogenic *PIK3CA* mutation (*H1047R* or *E545K*) to mutant KRAS is sufficient to convert cells from RAS-inhibitor-sensitive to largely resistant across several complementary models. This fits with earlier work in other *KRAS*-mutant cancers: in pancreatic cancer models, co-targeting PI3Kα or EGFR increased the activity of the KRAS-G12D inhibitor MRTX1133,^23^ and in pancreatic cancer patients treated with daraxonrasib itself, PI3K-pathway alterations were found in 9% of cases with acquired resistance^24^. Our work extends this concept to CRC and identifies *PIK3CA* mutation as a potential predictive biomarker of resistance rather than just an acquired resistance mechanism seen after treatment: patients already carrying a *PIK3CA* mutation at baseline may be less likely to respond to RAS inhibition, which could help guide treatment choice. Data from Choi et al.^25^ support this conclusion by demonstrating that response depended on how strongly each tumor relied on ERK versus PI3K signaling to maintain cyclin D1 levels. Our isogenic design, in which only the PIK3CA allele differs between clones, adds a complementary piece of evidence: changing PIK3CA genotype alone is enough to cause this resistance.

Having established that *PIK3CA* mutation drives resistance, we used our isogenic system to determine the effects of PI3K signaling in the context of KRAS inhibition. The classical function of PI3K/AKT/mTORC1 signaling is to control cap-dependent translation through 4E-BP1 phosphorylation. In our phosphoproteomic dataset, phosphorylation of 4E-BP1 at the canonical mTORC1 sites is increased in *PIK3CA*-mutant clones upon RAS(ON) inhibition. This confirms that the PI3K/mTORC1 axis remains specifically engaged in the mutant clones, but this translational signature alone is not the only candidate mechanism for the resistance phenotype. Instead, the dominant change we observe is a reduced apoptotic response to RMC-6236, together with a shift in phosphorylation of proteins involved in cell-cell adhesion and focal adhesion signaling (adherens junctions, PARVA, CRKL). Thus, in this co-mutated context, PI3K appears to act mainly as a survival and cell-remodeling signal via a blunting of apoptosis and a reorganization of adhesion/junction signaling, rather than primarily through its classical translational-control function. A recent study found that focal adhesion kinase (FAK) organizes PI3K-AKT signaling locally at focal adhesions, and that combined FAK and PI3Kα inhibition was more effective than either alone,^26^ supporting the idea that this focal-adhesion/survival signaling node, rather than translation, is the operative output of PI3K signaling here.

Our data show that pharmacologically combining KRAS and PIK3CA inhibition restores RMC-6236 efficacy in all of our co-mutated CRC models, through induction of apoptosis. A related combination strategy was effective in *KRAS-G12C*-mutant lung cancer models, where adding a selective mTORC1 inhibitor to a KRAS-G12C inhibitor produced synergistic inhibition of cyclin D1 and cap-dependent translation, together with upregulation of the pro-apoptotic protein BIM, leading to synergistic induction of apoptosis^27^. Given our finding that the PI3K role here is linked to focal-adhesion and survival signaling rather than translation, an alternative or complementary strategy worth considering is direct FAK inhibition instead of, or alongside, PIK3CA inhibition. In *KRAS-G12C*-mutant cancers, KRAS inhibition was shown to adaptively activate FAK, and combining a FAK inhibitor (IN10018) with KRAS-G12C inhibitors produced synergistic anti-tumor effects across multiple cell line, CDX, and PDX models by blocking FAK–YAP signaling^28^. Testing whether FAK inhibition can similarly resensitize our *PIK3CA*-mutant CRC models to RMC-6236 would help clarify whether PIK3CA mutation acts mainly through this focal-adhesion signaling node and could offer a therapeutic alternative with a different toxicity profile than direct PI3K inhibition.

A limitation of our study could be that the two PIK3CA alleles we tested (helical-domain E545K and kinase-domain H1047R) produced largely overlapping phosphoproteomic responses to RMC-6236, despite being activated through distinct biochemical mechanisms. Indeed, helical-domain mutants such as E545K remain dependent on binding to RAS-GTP, whereas the kinase-domain mutant H1047R is RAS-GTP-independent but instead requires binding to the p85 regulatory subunit for full catalytic activation^29^. Because our phosphoproteomic characterization was performed using a single representative CRISPR clone per genotype, we cannot fully exclude that this convergence partly reflects clone-specific effects rather than a true, generalizable property of each allele. This contrasts with findings in head and neck cancer cell lines, where different *PIK3CA* mutations were shown to build distinct protein interaction networks and confer different drug sensitivities, including a specific dependency on HER3 for some alleles^30^. It is possible that this reflects distinct effects of specific *PIK3CA* mutations in the context of KRAS vs. other oncogenic drivers. However, additional mechanistic work is required to assess this possibility.

As RMC-6236 and GDC-0077 (inavolisib) are both already FDA-approved drugs, this should make it easier to test their combination clinically although overlapping toxicity is a concern. However, allele-specific KRAS inhibitors seem far better tolerated and would likely be effective in combination with KRAS inhibition. Overall, although these data will need to be confirmed *in vivo*, they do support the clinical development of combined KRAS and PIK3CA targeting as a potential therapeutic strategy for KRAS/PIK3CA co-mutated CRC and identify PIK3CA mutation status as a candidate biomarker to select patients for this combination rather than RAS inhibitor monotherapy. They also identify *KRAS*-mutant CRC as an important context for the mechanistic study of PI 3-kinase activation and its downstream effects.

## Acknowledgements

We thank Drs. Kevan Shokat, Chloe Atreya, and JP Coppe and members of the Krogan and Gordan laboratories for their valuable feedback and discussion. This work was supported by NIH/NCI R37 (CA266549) funding and U54 funded NIH/NCI (CA274502) award.

## Supplementary Data

**Supplementary Data 1.**
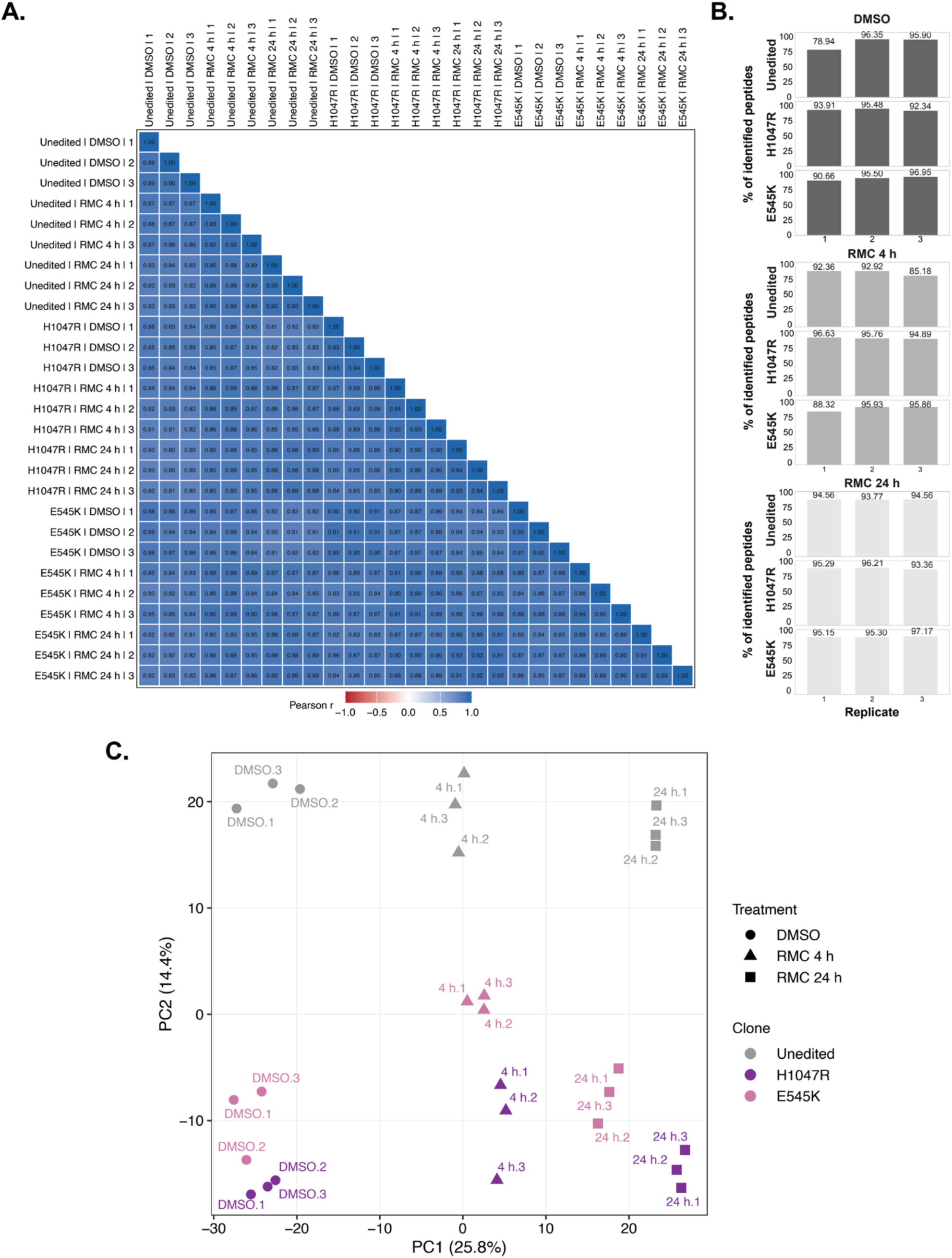
**related to Figure 3 and 4: A.** Pairwise Pearson correlation of log2 phosphosite intensities between the 27 LC-MS/MS runs, computed by artMS. Samples are ordered by clone and then by treatment. All 351 pairwise correlations lie between 0.80 and 0.94 (median 0.87), the color scale spans from −1 to +1. **B.** Phosphopeptide enrichment per run, given as the percentage of identified peptides carrying a phospho (STY)-modification. Phosphopeptides represent 78.9% to 97.2% of identified peptides (median 95.1%). Three runs fall below 90%, the lowest being the first unedited DMSO replicate at 78.9%. No sample was excluded from any analysis. **C.** Principal component analysis (PCA) of the 4,463 phosphosites quantified in all 27 runs without imputation. PC1 and PC2 account for 25.8% and 14.4% of the total variance. Color: clone; shape: treatment; labels: treatment and replicate number. PC1 orders the samples by treatment in all three clones, whereas PC2 separates the three clones from one another, including before treatment.

**Supplementary data 2.**
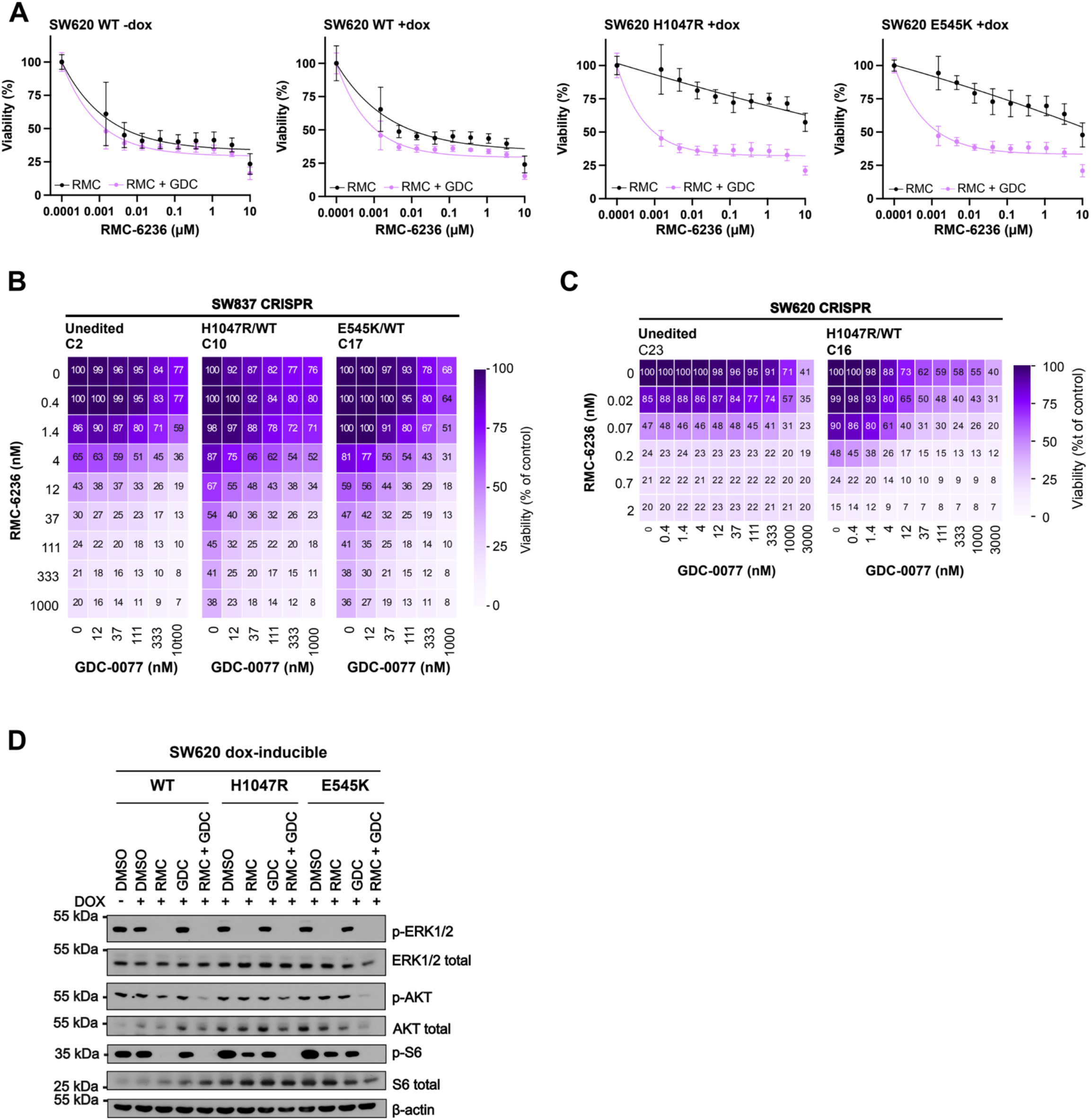
**related to figure 5. A.** SW620 FLAG-tagged, doxycycline-inducible PIK3CA (WT, H1047R, or E545K)-expressing cells were treated with increasing doses of RMC-6236, with or without GDC-0077 (200 nM), for 5 days and viability was assessed by CellTiter-Glo. WT −dox and +dox: uninduced and doxycycline-induced WT PIK3CA. **B.** SW837 CRISPR unedited (UE2), PIK3CA H1047R (clone 10), or PIK3CA E545K (clone 17) clones were treated with a dose matrix of RMC-6236 (0-1000 nM) and GDC-0077 (0-1000 nM) for 5 days and then subjected to CellTiter-Glo viability assay, a Matrix-response was computed. n = 3 independent experiments. **C**. SW620 CRISPR unedited (clone 23) or PIK3CA H1047R (clone 16) clones were treated with a dose matrix of RMC-6236 (0-1000 nM) and GDC-0077 (0-3000 nM) for 5 days and then subjected to CellTiter-Glo viability assay, a Matrix-response was computed. n = 3 independent experiments. **D.** Immunoblot of KRAS/MAPK and PI3K/AKT downstream signaling in response to RMC-6236 (100 nM) and GDC-0077 (200 nM), alone or in combination, for 24 h in SW620 dox-inducible cells. Loading control: β-actin.

**Supplementary data 3.**
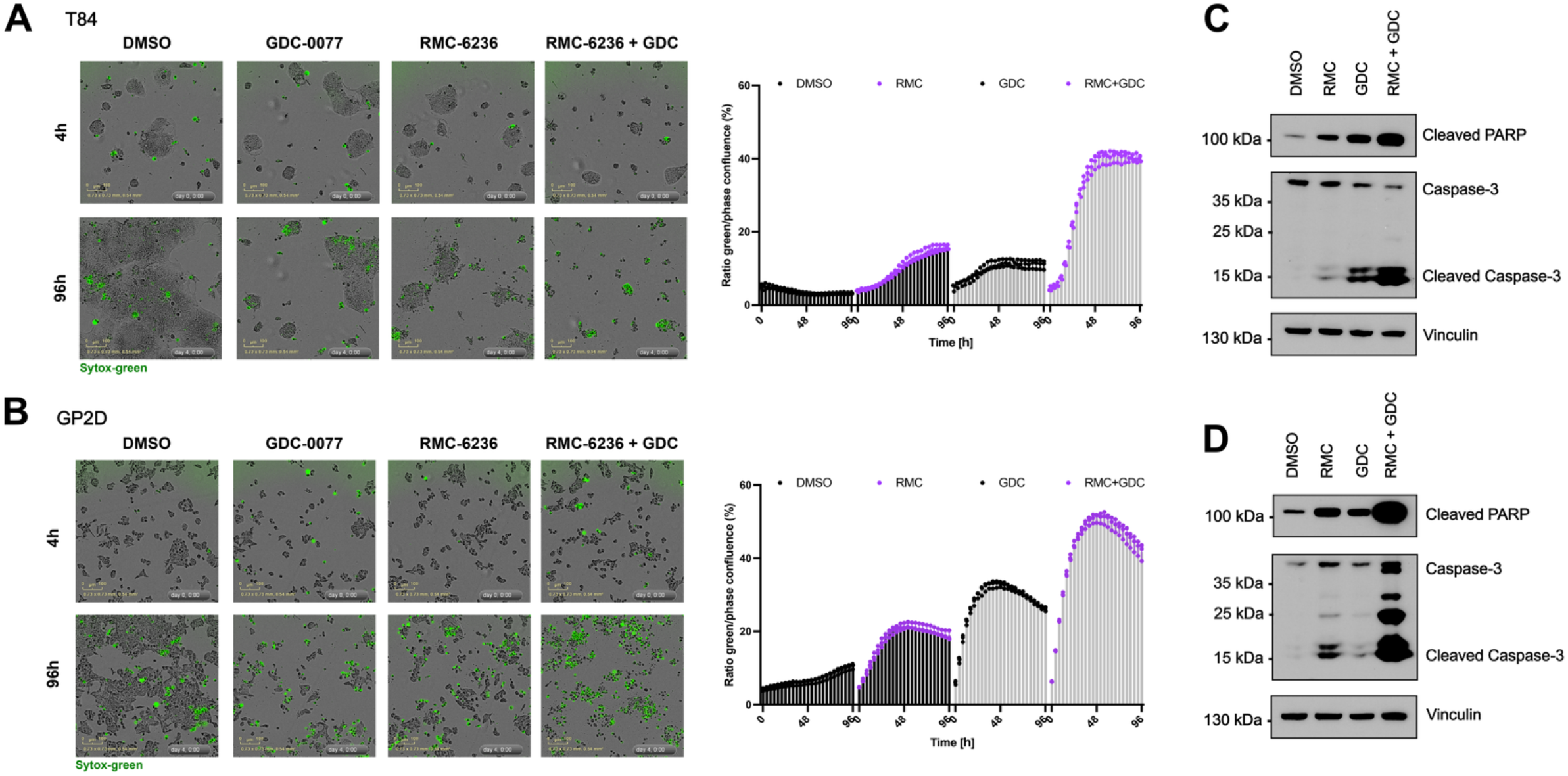
**related to figure 6. A-B.** SYTOX-cell death detection using live-cell microscopy in T84 (**A**) or GP2D (**B**) cells that were treated with RMC-6236 (500 nM) with or without GDC-0077 (500 nM) for 5 days. Representative images of each condition at 4 h and 96 h with phase and green fluorescence combined are shown. The ratio of Sytox-green positive cells over cell mask was quantified as a percentage in each condition. **C-D.** Western blot of apoptosis markers Cleaved PARP and Cleaved Caspase-3 in response to RMC-6236 (500 nM) and GDC-0077 (500 nM) combination for 24h in T84 (**C**) or GP2D (**D**).

## Bibliography

1. Simanshu, D.K., Nissley, D.V. & McCormick, F. RAS Proteins and Their Regulators in Human Disease. Cell 170, 17–33 (2017).

2. Hoadley, K.A. et al. Multiplatform analysis of 12 cancer types reveals molecular classification within and across tissues of origin. Cell 158, 929–944 (2014).

3. Cancer Genome Atlas Network Comprehensive molecular characterization of human colon and rectal cancer. Nature 487, 330–337 (2012).

4. Singhal, A., Li, B.T. & O’Reilly, E.M. Targeting KRAS in cancer. Nat Med 30, 969–983 (2024).

5. O’Reilly, E.M. et al. Daraxonrasib or Chemotherapy in Previously Treated Metastatic Pancreatic Cancer. N Engl J Med 395, 325–337 (2026).

6. Yaeger, R. et al. Adagrasib with or without Cetuximab in Colorectal Cancer with Mutated KRAS G12C. N Engl J Med 388, 44–54 (2023).

7. Fakih, M.G. et al. Sotorasib plus Panitumumab in Refractory Colorectal Cancer with Mutated KRAS G12C. N Engl J Med 389, 2125–2139 (2023).

8. Carracedo, A. et al. Inhibition of mTORC1 leads to MAPK pathway activation through a PI3K-dependent feedback loop in human cancer. J Clin Invest 118, 3065–3074 (2008).

9. Castellano, E. & Downward, J. RAS Interaction with PI3K: More Than Just Another Effector Pathway. Genes Cancer 2, 261–274 (2011).

10. Hsiao, Y. et al. Analysis and visualization of quantitative proteomics data using FragPipe-Analyst. bioRxiv (2024).

11. Yu, F., Haynes, S.E. & Nesvizhskii, A.I. IonQuant Enables Accurate and Sensitive Label-Free Quantification With FDR-Controlled Match-Between-Runs. Mol Cell Proteomics 20, 100077 (2021).

12. Shteynberg, D.D. et al. PTMProphet: Fast and Accurate Mass Modification Localization for the Trans-Proteomic Pipeline. J Proteome Res 18, 4262–4272 (2019).

13. Casado, P. et al. Kinase-substrate enrichment analysis provides insights into the heterogeneity of signaling pathway activation in leukemia cells. Sci Signal 6, rs6 (2013).

14. Wiredja, D.D., Koyuturk, M. & Chance, M.R. The KSEA App: a web-based tool for kinase activity inference from quantitative phosphoproteomics. Bioinformatics 33, 3489–3491 (2017).

15. Gingras, A.C. et al. Hierarchical phosphorylation of the translation inhibitor 4E-BP1. Genes Dev 15, 2852–2864 (2001).

16. Huang, R. et al. The NCATS BioPlanet - An Integrated Platform for Exploring the Universe of Cellular Signaling Pathways for Toxicology, Systems Biology, and Chemical Genomics. Front Pharmacol 10, 445 (2019).

17. Song, K.W. et al. RTK-Dependent Inducible Degradation of Mutant PI3Kalpha Drives GDC-0077 (Inavolisib) Efficacy. Cancer Discov 12, 204–219 (2022).

18. Kavgaci, G. et al. Evaluation of KRAS and NRAS mutations in metastatic colorectal cancer: an 8-year study of 10 754 patients in Turkey. Mol Oncol 19, 2967–2977 (2025).

19. Zha, J., Harada, H., Yang, E., Jockel, J. & Korsmeyer, S.J. Serine phosphorylation of death agonist BAD in response to survival factor results in binding to 14-3-3 not BCL-X(L). Cell 87, 619–628 (1996).

20. Datta, S.R. et al. Akt phosphorylation of BAD couples survival signals to the cell-intrinsic death machinery. Cell 91, 231–241 (1997).

21. Brunet, A. et al. Akt promotes cell survival by phosphorylating and inhibiting a Forkhead transcription factor. Cell 96, 857–868 (1999).

22. Ianevski, A., Giri, A.K. & Aittokallio, T. SynergyFinder 2.0: visual analytics of multi-drug combination synergies. Nucleic Acids Res 48, W488–W493 (2020).

23. Hallin, J. et al. Anti-tumor efficacy of a potent and selective non-covalent KRAS(G12D) inhibitor. Nat Med 28, 2171–2182 (2022).

24. Aronchik, I. et al. Acquired resistance to the RAS(ON) multi-selective inhibitor daraxonrasib guides rational combination therapy strategies in pancreatic cancer. Nat Med 32, 2865–2877 (2026).

25. Choi, P.H. et al. Distinct ERK and PI3K dependencies mediate a heterogeneous response to RAS(ON) inhibition in KRAS-mutant colorectal cancer. Cell Rep 45, 117559 (2026).

26. Wang, J. et al. Spatial organization of PI3K-PI(3,4,5)P(3)-AKT signaling by focal adhesions. Mol Cell 84, 4401–4418 e4409 (2024).

27. Kitai, H. et al. Combined inhibition of KRAS(G12C) and mTORC1 kinase is synergistic in non-small cell lung cancer. Nat Commun 15, 6076 (2024).

28. Zhang, B. et al. Focal Adhesion Kinase (FAK) Inhibition Synergizes with KRAS G12C Inhibitors in Treating Cancer through the Regulation of the FAK-YAP Signaling. Adv Sci (Weinh) 8, e2100250 (2021).

29. Zhao, L. & Vogt, P.K. Helical domain and kinase domain mutations in p110alpha of phosphatidylinositol 3-kinase induce gain of function by different mechanisms. Proc Natl Acad Sci U S A 105, 2652–2657 (2008).

30. Swaney, D.L. et al. A protein network map of head and neck cancer reveals PIK3CA mutant drug sensitivity. Science 374, eabf2911 (2021).

